# Drug repurposing for yellow fever using high content screening

**DOI:** 10.1101/225656

**Authors:** Denise R. B. Pilger, Carolina B. Moraes, Laura H. V. Gil, Lucio H. Freitas-Junior

## Abstract

**Background:** Yellow fever is an acute febrile illness caused by a mosquito-borne flavivirus. The discovery that mosquitos were responsible for transmission and that the disease was preventable by vector control, as well as the development of an efficacious vaccine (in the 1930s) have reduced its public health impact. However, underutilization of vaccination and discontinuation of vector control measures have led to yellow fever outbreaks, affecting thousands of people in Africa and South America, and is a continued threat to people who travel to endemic regions without vaccination. An additional concern is that the vaccine has important contraindications, and the currently available doses are insufficient to immunize the populations under risk of contracting yellow fever. Specific antiviral chemotherapy would be an alternative to vaccination, however there is none yet available. Drug repurposing is one strategy that can speed up drug discovery and development and has been underexplored for yellow fever.

**Methods:** A high content screening assay for the discovery of anti-yellow fever compounds was developed and used to screen a library of pharmacologically compounds, which includes approved drugs and covers most signalling pathways and all major drug target classes. The same library was screened against cells infected with a serotype 2 of the dengue virus.

**Results and conclusion:** From 1280 compounds screened, 88 compounds (6,9% of the library) were found to reduce yellow fever virus (YFV) infection in 50% or more. Interestingly, the number of compounds that presented similar activity against dengue (DENV) infection was considerably lower (18 compounds, 1,4%). Among these, 12 compounds were active against both viruses, highlighting the potential of finding compounds with broad antiviral activity in diversity libraries. The top 27 anti-YFV compounds were selected for further activity determination in dose-response. Five compounds (Gant61, benztropine mesylate, brequinar sodium salt hydrate, PF-429242 dihydrochloride and U-73343) presented selective activity against YFV infection in a human cell line, but only two of them, brequinar and U-73343, were also active against DENV. These compounds, which represent a broad spectrum of pharmacological functions, have not been previously described as having anti-YFV activity, thus offering a valuable opportunity for the development of specific antiviral chemotherapy for yellow fever.

## INTRODUCTION

Flavivirus is a large genus comprising more than 70 pathogenic viruses of different pathologies with significant public health impact in across the tropical world. Yellow fever virus (YFV) and dengue virus (DENV) are some of the most important human pathogenic flaviviruses, sharing the same vectors and causing similar symptoms, ranging from low fever and malaise in mild infections, to haemorrhage and rapid terminal events with shock and failure of various organs in haemorrhagic fever syndrome^1,2^.

Yellow fever is endemic in several South American and African countries. It has been estimated that up to 1.7 million yellow fever cases occur in Africa each year, mostly in West African countries, resulting in 29,000–60,000 deaths^3,4^. Since late 2016, yellow fever outbreaks have occurred in the Southeast region of Brazil, with 792 confirmed cases, including 274 deaths^5,6^ in about 12 months. The transmission of YFV occurs through the bite of mosquito vectors belonging to different species. Usually sporadic human infection occurs through occasional transmission of YFV from nonhuman primates to humans through mosquito vectors belonging to the sylvatic YFV cycle – from the *Haemagogus* and *Sabethes* genus in South America and the *Aedes* genus in Africa – when humans come into endemic jungle and forest areas^7,8^. In the urban cycle, the ubiquitous vector *Aedes aegypti* is responsible for the human-to-human transmission. The disease can spread rapidly and affect thousands of individuals^8,9^.

Yellow fever is often an acute disease with a range of clinical manifestations, including fever, bradycardia and leukopenia. In 25 to 50% of cases, the acute phase can progress to an intoxication stage characterized by high fever, haemorrhagic syndrome, jaundice and kidney disease, which can be fatal. The patients surviving the acute period enter the convalescence phase, characterized by prolonged weakness and fatigue that can last several weeks ^3,4,10^. Thus, in addition to presenting high lethality rates, yellow fever also causes heavy socio-economic burden in afflicted populations.

The adoption of vector control measures in the early 20^th^ century, followed by the development of a very effective vaccine in the 1930s, have greatly reduced yellow fever occurrence and deaths over the past decades^4^. However, the yellow fever vaccine also presents important contraindications, as it may cause severe adverse effects, and current protocols do not recommend the yellow fever vaccine to be administered to elderly older than 60 years, pregnant and breastfeeding women and immunocompromised individuals^11^. Another issue is the vaccine production worldwide, as it currently amounts to 80 million doses/year, whereas there are more than 3 billion people currently living in risk areas where *Aedes* mosquitos are endemic, offering the potential for yellow fever (re)introduction^12,13^. Furthermore, deforestation and poor urbanization, present in most endemic countries, combined with changes in global migration patterns, have created ideal conditions for outbreaks of urban yellow fever in previously disease-free areas^7^.

Given the current yellow fever scenario, a specific antiviral treatment would have considerable value for populations that cannot take the vaccine. Moreover, in the wake of an outbreak, an antiviral could be administered either as prophylaxis or at the onset of any symptoms that could be associated with the disease, as strong immunity (80% of protection) normally needs at least 10 days after vaccination to develop^11,14^. However, there is still no antiviral or other specific chemotherapy for the treatment of YFV, and clinical management of disease involves only attenuation of symptoms and supportive care of patients^15^.

Despite the need, there are very few published studies of drug screening for yellow fever. One feasible and often-deployed strategy for drug discovery is drug repurposing or repositioning, which consists on finding new uses for known clinical and experimental drugs. As several pharmacological, toxicity and formulation aspects are already known for these molecules, they can be more readily available for clinical trials when compared to novel drugs.

Here we report the development of the first high content screening assay for the discovery of inhibitors of YFV infection. This assay was used to screen a compound library of pharmacologically active compounds, including pre-clinical and clinical drugs. The same library was screened against dengue virus in an analogous assay, enabling the comparison of results and the search for broad-spectrum antiflaviviral drugs.

## METHODOLOGY

### Virus and cell culture

A yellow Fever viral strains 17D expressing the enhanced yellow fluorescent protein (eYFP) was constructed for the HCS assay. The *Aedes albopictus* C6/36 cell line, the human hepatome cell line Huh7 and dengue virus serotype 2 isolate (DENV2) were kindly provided by Dr. Amílcar Tanuri from Federal University of Rio de Janeiro. C6/36 cells were cultivated in Leibovitz L-15 media (L4386, Sigma-Aldrich) at 28 °C. Huh7 cells were cultivated in DMEM F-12 media (56498C, Sigma-Aldrich) at 37 °C, 5% CO_2_. In both cases the media was supplemented with 10% FBS (F0804, Sigma-Aldrich), 100 U/mL of Penicillin and 100 μg/mL of Streptomycin (P4333, Sigma-Aldrich). Viral propagation and quantification was performed as described^16^ with exception of the viral titration, which was carried out in Huh7 cells.

### Construction and recovery of YFV expressing the eYFP gene

A recombinant YFV expressing enhanced yellow fluorescent protein (eYFP) was constructed by homologous recombination in yeast, using the previously constructed pBSC-YFV17D-T7 plasmid, which contains the entire genome of YFV strain 17D. The insertion of eYFP into YFV genome was made according to the strategy described by Bonaldo et al, 2007^17^, who introduced the GFP gene between the E and NS1 genes of YFV strain 17D. The YFV-eYFP plasmid was assembly by yeast recombination of two PCR products and the Nar-I digested pBSC-YFV17D-T7 vector. The first PCR fragment (808 base pairs) was amplified with forward primer (5’GATGTTTTTGTCTCTAGGAGTTGGGGCGGACCAGGGCTGTGCAATTAATTTCGGGGGCGCCATGGTGAGCAAGGGCGAGG) that contained homologous sequences for recombination with the E gene, plus the first 27 nucleotides of the NS1 gene, Nar-I site sequence and 19 first nucleotides from eYFP gene. The reverse primer (5’-TCGGGCGGTTGCTTCGAACATTTTGGCGCCCTTGTACAGCTCGTCCATGC) contained the first 20 nucleotides of the DENV1 stem-anchor region for recombination, followed by a Nar-I restriction site sequence and 20 first nucleotides of eYFP gene. The second PCR fragment (754 base pairs) was amplified with the following primers pair: Forward (5’-GGCGCCAAAATGTTCGAAGCAACCGCC) and Reverse (5’-GGAAAGAATTCCAGACC CGG). This PCR product contained homologous sequence for recombination with the first PCR product (terminal eYFP gene sequence), followed by DENV1 stem-anchor region sequence (288bp, position 2132 to 2819; GeneBank accession number JX669474) and the sequence for recombination in NS1 gene. The generated pBSC-YFV-eYFP plasmid was used as a template for in vitro RNA synthesis (Megascript, Ambion). RNA transcripts were electroporated into BHK-21 cells and virus replication was verified by indirect immunofluorescence e reporter gene expression, as described previously^18^.

### Reference drug and compound library preparation

The reference drug Interferon α 2A (IFNα 2A) and the Library of Pharmacologically Active Compounds (LOPAC^1280^) were purchased from Sigma-Aldrich. IFNα 2A was diluted in Dulbecco’s Phosphate-buffered saline (DPBS) containing 0.5% (W/V) bovine serum albumin (SH30574.03, HyClone) to a final concentration of 5.2 nM. The compound library was prepared as 1 mM stock solutions in 100% DMSO and arrayed in 384-well stock plates.

### High content screening assay

Before each assay, 1.2 μL of library compound solution was transferred from stock plates onto polypropylene 384-well plates (Greiner Bio-One, Cat. No. 784201), containing 18.8 μL of DPBS pH 7.4, yielding a final concentration of 600 μM compound in 6% of DMSO (v/v, 16,67-fold dilution). Then, 10 μL of compound solution were transferred into tissue culture-treated black polystyrene 384-well assay plates (Greiner Bio-One, Cat. No. 781091). Following compound transfer, 50 μL/well of a mixture of Huh7 cells at 4x10^4^ cells/mL and YFV viral particles at a multiplicity of infection (MOI) of 2.5 were added in each well. The final LOPAC compound concentration was 10 μM in 1% DMSO (v/v) in all wells. For the DENV2 assay, 50 μL/well of a mixture of Huh7 cells at 6x10^4^ cells/mL and DENV2 at a MOI of 0.5 were added in each well. All plates included vehicle (DMSO)-treated YFV or DENV2-infected cells as negative controls, and non-infected (mock-infected) as well as and YFV or DENV2-infected cells treated with 5.2 nM IFNα 2A as positive controls. The plates were incubated in a humidified atmosphere at 37 °C, 5% CO_2_, followed by fixation with 4% paraformaldehyde in DPBS pH 7.4 after 72 h. YFV plates were stained with 5 μg/mL of 4′,6-Diamidine-2′-phenylindole dihydrochloride (DAPI). Cells infected with DENV2 were submitted to indirect staining of viral infection by immunofluorescence as described previously^19^. Briefly, the D1-4G2-4-15 monoclonal antibody, which recognizes the E protein from flavivirus, was used as primary antibody, and the goat anti-mouse IgG (H+L) conjugated with AlexaFluor594 (A11032, Thermo), as a secondary antibody. Four images from each well were acquired using the Operetta High Content Imaging System (Perkin Elmer) with a 20X magnification objective.

### Data analysis

All images were analyzed and processed with the High Content Analysis Software Harmony (Perkin Elmer), which is able to detect host cells as well as YFV in the cell cytoplasm through the eYFP signal and DENV2 through the AlexaFluor594 signal. The software provides as output data from each well the total number of host cells, number of infected cells, rate of infection per well and signal strength of eYFP inside the cell cytoplasm. The cell ratio (CR) was defined as the number of cells in the tested compound well divided by the mean number cells in negative control wells, while the infection ratio (IR) was defined as the total number of infected cells in all images from the well divided by the total number of cells in all images from the same well. The raw data for IR values were normalized to negative (non-treated infected cells) and positive controls (infected cells treated with Interferon α 2A) to determine the normalized antiviral activity, according to the equation below, as described elsewhere^20^. The correlation coefficient (*r*) of the two YFV screening runs was calculated using Pearson correlation in GraphPad Prism software, version 6.

### Hit selection and activity confirmation in dose-response

Following exclusion of cytotoxic compounds, the top 27 most active compounds for YFV screen were selected as hits for confirmatory screening in dose-response. These compounds were cherry-picked and serially diluted in a 1:2 ratio in DMSO (v/v) to produce 10-concentration test points, starting at 100 μM, with exception of Benztropine mesylate, which started at 12.5 μM. Dose-response curves were calculated using the sigmoidal dose-response – variable slope function from the GraphPad Prism version 6 with the default settings. The EC_50_ was defined as the effective concentration resulting in 50% inhibition of YFV or DENV infection, and maximum activity values were determined from fitted curves from normalized activity datasets. The CC_50_ value, defined as the compound concentration resulting in a 50% reduction in cell number compared with the positive control (non-infected cells), was used to evaluate cell toxicity. The Selectivity Index (SI) was calculated as SI = CC_50_/EC_50_.

## RESULTS

### Activity assay development with Yellow Fever Virus

The yellow fever 17D vaccinal strain expressing the yellow fluorescent protein (YFV-eYFP), constructed as depicted in Figure 1A, was adapted to a high content screening assay for assessment of antiviral activity of compounds. Four images were acquired from all conditions and processed with the High Content Analysis Software Harmony to determine the total number of cells, number of infected cells, ratio of infected cells, and mean signal of eYFP/AlexaFluor594 (Figure S1). In preliminary studies, the optimal number of host cells per well, DMSO concentration, multiplicity of infection and the assay duration were determined (Figure S2 and S3). The DENV2 assay was adapted, with minor changes, from the previously published methodology^19^. Huh7 cells grow as a monolayer and present a large cytoplasm that allow for improved quantification of viral infection intensity in the host cell (Figure 1B). The reference drug IFNα 2A was chosen for these assay, as it was previously reported to have broad spectrum anti-flaviviral activity^21^. The IFNα 2A antiviral activity and cell toxicities against YFV and DENV2 infection of Huh7 were determined by dose-response curves. Figure 1C shows the dose-response curves of the reference compound, with the percent normalized activity and cell ratio. It was observed that IFNα 2A showed inhibition of YFV and DENV2 infection in a dose-dependent manner. The reference compound did not present cytotoxicity even at very high concentration (5.2 nM). The EC_50_ values obtained for YFV and DENV2 were 17.31 pM and 30.07 pM, respectively.

**Figure 1.**
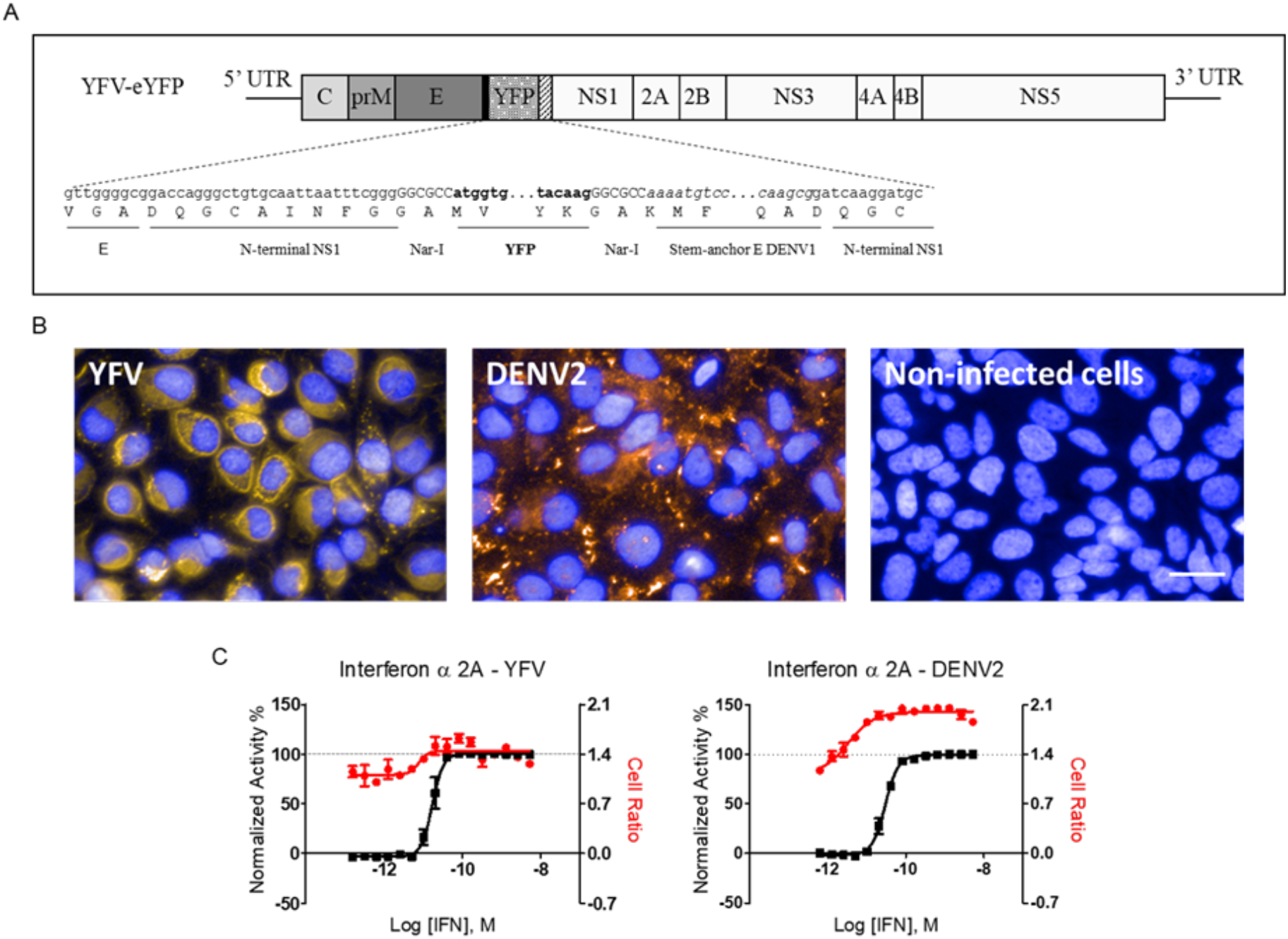
Development of High Content Screening assay for yellow fever. **A)** Genome organization of YFV-eYFP shows the eYFP cloned between E and NS1 genes. The nucleotides and amino acid sequences (single-letter code) of the E (C-terminus region), N-terminal NS1 (first 27 nucleotides), stem-anchor E of DENV-1 (partial), eYFP reporter gene (bold) and N-terminal NS1 boundary are indicated. The eYFP is flanked on both sites by Nar-I recognition restriction sites (uppercase). **B)** Representative images showing YFY-infected (left), DENV2-infected (middle) and non-infected Huh7 cells. The cell nuclei were stained with DAPI (blue), YFV represented by eYFP signal (yellow) and DENV2 infection detected by immunofluorescence (orange); bar, 40 μm. **C)** Dose-response curves of the reference drug Interferon α 2A against yellow fever (left) and dengue virus-infected cells (right). The X-axis indicates the log of Interferon α 2A concentration (molar), the right Y-axis shows the normalized antiviral activity in percentage (in black), which represents the inhibition of infection in relation to non-infected control, and the right Y-axis shows the cell ratio (in red). Data points show the mean (squares) and standard deviation (bars) values from two independent assays.

### Screening of the LOPAC^1280^ to identify drugs that are effective against Yellow Fever virus and Dengue virus serotype 2

The YFV HCS assay was validated by means of screening the Library of Pharmacologically Active Compounds (LOPAC), containing 1280 compounds, at 10 μM. The library was screened in two independent runs, and the correlation coefficient (*r*) between the two runs was 0.85 for compounds and controls and 0.80 for compounds only (Figure 2A and Figure S4). The mean Z’-factor of the runs were 0.78 ± 0.14 (mean ± standard deviation) for run 1, and 0.92 ± 0.04 for run 2. The DENV2 HCS assay had been previously validated (data not shown), thus the LOPAC was screened a single time, also at 10 μM. The average Z’-factor for all plates was 0.97 ± 0.01 (mean ± standard deviation).

**Figure 2.**
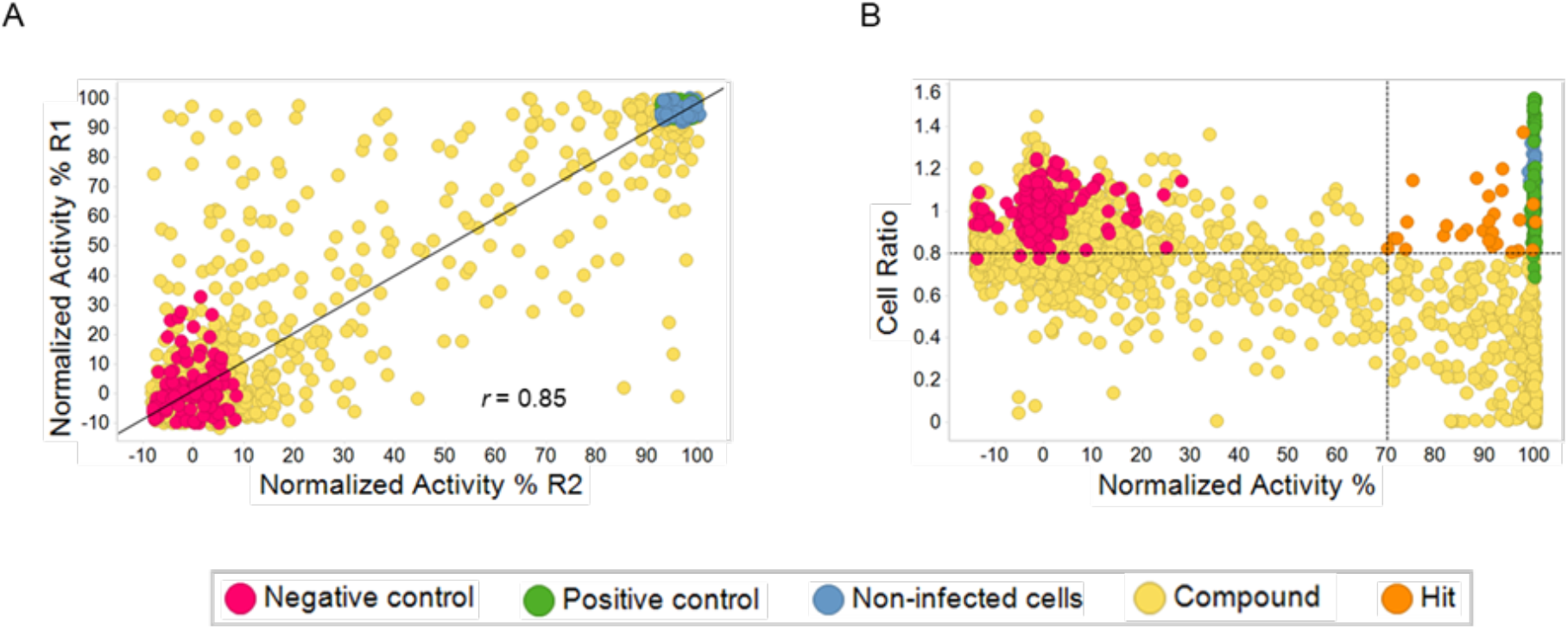
Validation of the YFV HCS assays with the LOPAC^1280^ library. **A)** Correlation the normalized activity, with and the correlation coefficient (*r*) of the two runs. Dots represent treated wells from the two independent replicates. **B)** Scatter-plot distribution of compounds according to antiviral activity (X-axis) and host cell toxicity (Y-axis). Each spot represents one well from the two independent replicates. Dotted lines indicate the cut-off values of the primary hits based on the criteria: > 70% normalized activity and > 0.8 cell ratio, to select the top 27 hits.

As the YFV and DENV assays are similar (both use the same host cell and last for the same period of viral infection and compound exposure) and the screening in both deployed the same library, at the same concentration, a comparison of antiviral activity was undertaken in search for compounds that might have activity against both viruses. The compounds with antiviral activity > 50% and cell ratio > 0.5 were compared from both YFV and DENV screening. In the YFV screening, there were 88 compounds in this category, while this number was reduced to 18 compounds in the DENV screening (Figure S5). Among those, only 12 compounds were in common against both viruses (data not shown). Altogether, the data highlight that anti-DENV are more difficult to obtain than anti-YFV hits, however, most compounds that display anti-YFV activity were also at least partially active against DENV2 (12 out of 18 hits), demonstrating that there is a good potential for the discovery of compounds with broad antiviral activity with this type of approach.

The top 27 YFV screening hits yielded normalized activity equal to or approximately 70% and host cell toxicity ≤ 20 (Figure 2B) and were selected for activity confirmation in dose-response. The twenty-seven YFV primary hits were tested in dose-response, and 25 of them presented high antiviral activity. However, most presented cell toxicity at the higher concentrations (Table S1). To be progressed further into drug discovery, a screening hit compound should have an acceptable *in vitro* potency (usually EC_50_ values < 10 μM) and sufficient selectivity toward the pathogen as opposed to mammalian cells (SI ≥ 10)^22^, among other criteria. Among the confirmed YFV selected compounds, five showed a hit-like activity profile: Gant61 (Pubchem CID: 421610), Benztropine Mesylate (Pubchem CID: 238053), Brequinar sodium salt hydrate (Pubchem CID: 86346664), PF-429242 dihydrochloride (Pubchem CID: 71296088), and U-73343 (Pubchem CID: 114825). From these five compounds, only two, Brequinar and U-73343, showed inhibition against both YFV and DENV2 infection. Brequinar presented the same level of inhibition of infection (normalized activity of approximately 98%) at 10 μM against both viruses. The compound U-73343 exhibited higher antiviral activity against YFV (100%) than against DENV2 (83%, data not shown). Both compounds displayed little to no cytotoxicity under the assays conditions, suggesting that both have potential to be further investigated as broad-spectrum antiflaviviral compounds.

With exception of Brequinar, none of these compounds were previously reported as active against Yellow Fever virus, although they had all been reported as potential antiviral against other viruses, including at least one other flavivirus (Table 1). Altogether, these data demonstrate these compounds have a good potential of being developed further as broad-spectrum anti-flavivirals, and may represent valuable starting points for anti-yellow fever drug discovery programs.

**Table 1.**
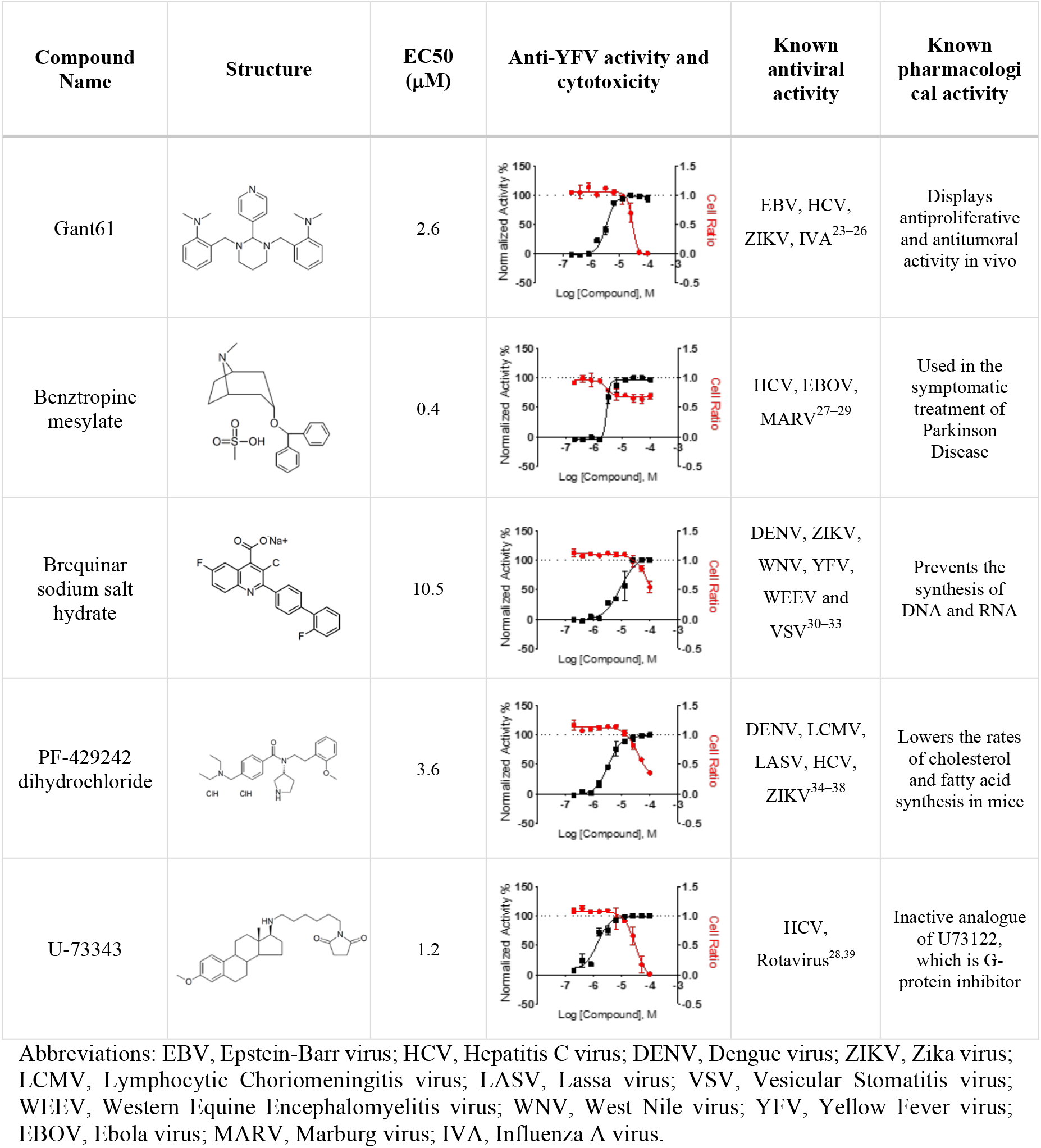
Molecular structure, antiviral activity and pharmacological activity of the anti-YFV with hit-like activity profile.

| Compound Name | Structure | EC50 (μM) | Anti-YFV activity and cytotoxicity | Known antiviral activity | Known pharmacological activity |
| --- | --- | --- | --- | --- | --- |
| Gant61 |  | 2.6 |  | EBV, HCV, ZIKV, IVA<sup>23–26</sup> | Displays antiproliferative and antitumoral activity in vivo |
| Benztropine mesylate |  | 0.4 |  | HCV, EBOV, MARV<sup>27–29</sup> | Used in the symptomatic treatment of Parkinson Disease |
| Brequinar sodium salt hydrate |  | 10.5 |  | DENV, ZIKV, WNV, YFV, WEEV and VSV<sup>30–33</sup> | Prevents the synthesis of DNA and RNA |
| PF-429242 dihydrochloride |  | 3.6 |  | DENV, LCMV, LASV, HCV, ZIKV<sup>34–38</sup> | Lowers the rates of cholesterol and fatty acid synthesis in mice |
| U-73343 |  | 1.2 |  | HCV, Rotavirus<sup>28,39</sup> | Inactive analogue of U73122, which is G-protein inhibitor |

Abbreviations: EBV, Epstein-Barr virus; HCV, Hepatitis C virus; DENV, Dengue virus; ZIKV, Zika virus; LCMV, Lymphocytic Choriomeningitis virus; LASV, Lassa virus; VSV, Vesicular Stomatitis virus; WEEV, Western Equine Encephalomyelitis virus; WNV, West Nile virus; YFV, Yellow Fever virus; EBOV, Ebola virus; MARV, Marburg virus; IVA, Influenza A virus.

## DISCUSSION

Despite the severity of the diseases and the clear need for chemotherapy, the body of published research on antiviral agents for yellow fever is less substantial in comparison with the amount of work undertaken for other major arthropod-borne flaviviruses, such as dengue. The first screening campaign to identify potential candidates with activity against YFV utilized a replicon-based HTS assay, which was used to screen 34,000 small molecule compounds and resulted in the discovery of two compounds (CCG-4088 and CCG-3394) that were active against YFV^40^. However, their antiviral efficacies in animal models have not been reported so far. Another campaign used dengue virus for a pilot screening of a small molecule library of 26,900 compounds. The screening resulted in 14 compounds that inhibit dengue virus replication, and only one, benzothiazolylphenyl urea, was selected to further studies. This compound also inhibited YFV replication, and animal studies showed that it was well-tolerated and protected 90% of YFV-infected hamsters from death, significantly reduced the viral load, and normalized liver function^41,42^. High Content Screening, which utilizes automated microscopy and image analysis, is another cell-based strategy for the discovery of antivirals and was adapted in this work. HCS assays involve the treatment of cell lines infected with viruses, replicons or pseudoinfectious particles (PIPs), with small molecule or natural product compounds. In all cases, the selection of hits is based on the reduction of infection measured by a fluorescent reporter gene or by indirect immunofluorescence^19,20,43^.

In the present work, the HCS assay developed covers the whole viral cycle, from entry, RNA synthesis and viral egress of the host cell, potentially exposing all druggable targets, including host molecules that participate in the viral infection process. We selected the top 20 compounds for each virus among the 1280 compounds, although, interestingly, there was little overlap in terms of antiviral activity between the top 20 most active compounds from each screen, with only two compounds presenting activity against both viruses. The inability of YFV hits to inhibit DENV2 infection (and vice-versa) can be attributed to intrinsic assay differences, of which the more likely are structural and functional differences of viral proteins, or differential use of host proteins and machinery, such as receptors and other proteins involved in signalling pathways. Conversely, the two compounds that showed antiviral activity in both screenings, U-73343 and Brequinar, are more likely to act on conserved viral proteins that regulate viral replication and assembly, such as NS3 protease and NS5 polymerase. These compounds have been previously described as having antiviral activity in other models, including HCV, another flavivirus, demonstrating their potential to be further explored as broad-spectrum antivirals.

Of the 27 selected anti-YFV compounds for activity confirmation in dose-response, only five were found to be sufficiently selective (ie., at least 10 times more active toward the virus than to the host cell) to merit further analysis of their potential as hits for optimization. All these five selected compounds (Gant61, Benztropine mesylate, Brequinar, PF-429242 and U-73343) were previously describes as having antiviral activity against several viruses, including HCV, DENV2 and even YFV (in the case of Brequinar), further validating the HCS approach as a viable technology for the discovery of antivirals.

Gant61 inhibits the Hedgehog (HH) pathway by inhibiting the transcriptional activity of GLI1/2. The HH pathway regulates critical cell decisions, including proliferation and apoptosis and also modulates wound healing responses in liver tissue^44,45^. A study found that higher levels of HH pathway activity was associated with cirrhosis and human hepatocellular carcinomas in patients with chronical viral hepatitis, and that the pathway inhibition reduced cell fibrogenesis, angiogenesis, and growth^46^. The treatment of HCV-infected primary fibroblast with Gant61 was able to counteract the fibrosis and also reduce HCV replication ^23,46^. This pathway is also correlated with the pathological outcome of the severe form of the yellow fever, which includes direct viral cytopathic effect, necrosis and apoptosis of hepatocyte cells. A therapeutic approach that blocks the mechanisms that trigger apoptosis, and in thesis, would decrease the pathogenesis of yellow fever, could protect the liver in severe cases of disease^47,48^. The reduction of YFV replication in our assay may be correlated with the apoptosis regulated through the HH pathway. However, Gant61 is still an experimental drug and more preclinical and clinical trials are necessary to understand its pharmacokinetics and toxicity^49^.

Benztropine mesylate is a muscarinic receptor antagonist used for symptomatic treatment of Parkinson’s disease. This compound has been demonstrated to have antiviral activity against HCV. One study demonstrated that benztropine inhibits HCV infection by blocking an early event, downstream of virion attachment but preceding HCV RNA replication^50^. Another study reported that benztropine inhibits Marburg and Ebola virus entry through different classes of G protein-coupled receptors (GPCRs), including the muscarinic acetylcholine receptor ^27^. It is known that different viruses use different strategies to hijack the GPCRs and the GPCR-activated signalling pathways. Although the entry through GPCR is not extensively described for flavivirus, Le Sommer et al. 2012^51^ reported that two members of the G protein-coupled receptor kinase family (GRK2 and GRK4) are required for efficient propagation of flaviviruses and that a decrease in GRK2 level alters both virus entry and RNA genome amplification. As benztropine mesylate is an antagonist of the muscarinic receptor, a member of the GPCRs family, it can be speculated that this compound inhibits YFV entry by antagonizing muscarinic receptors.

Brequinar sodium salt hydrate has broad-spectrum antiviral activity against both positive- and negative-strand RNA viruses. Brequinar inhibits cellular dihydroxyorotate dehydrogenase (DHODH), the fourth enzyme in the *de novo* pyrimidine biosynthesis pathway, preventing the synthesis of uracil nucleotides, and, consequently, DNA and RNA synthesis. The main antiviral mechanism of Brequinar is the depletion of intracellular pyrimidine triphosphate, thus impairing viral RNA replication^31,33^. Long Yeo et al. 2015^52^ found that Brequinar can potentiate interferon-stimulated response element (ISRE) activation induced by interferon. This result indicate that Brequinar can enhance exogenous IFN-induced ISRE activation. However, the *in vitro* efficacy did not translate into *in vivo* efficacy, because uridine uptake from diets replenish or maintain a high concentration of pyrimidine in the plasma, and therefore lowering the compound-mediated inhibition of viral replication^53^. Another problem is that resistance can develop, and could occur via various mechanisms^32^. Qing et al. 2010^33^ shown that Brequinar has activity against a broad spectrum of viruses by reducing the RNA synthesis, but dengue virus developed resistance through mutations in the envelope protein or NS5 gene. So far, it was not approved for clinical use for viral diseases owing to a low therapeutic index^32^.

Subtilisin kexin isozyme-1/site-1 protease (SKI-1/S1P), a key regulator of the cholesterol and fatty acid and their metabolism, through the processing of the transcription factor sterol regulatory element-binding proteins (SREBPs). It also regulates endoplasmic reticulum (ER)-stress response through cleavage of ATF6 and is involved in the processing of probrain-derived neurotrophic factor ^36,54^. The amino-pyrrolidine amide compound PF-429242 efficiently blocked the activation of SREBP by blocking the SKI-1/S1P-mediated process in CHO, HepG2 and HEK293 cell cultures and in CD1 mice^55^. Pasquato et al. 2012^35^ demonstrated that PF-429242 was able to block the biosynthesis of fusion-active mature glycoprotein precursor of the old-world arenaviruses LCMV and LASV, and showed potent anti-viral activity against LCMV and LASV in acute infection in cultured cells. They also demonstrated that the infection did not re-emerge after the interruption of drug treatment, indicating that PF-429242 treatment resulted in virus extinction. DENV utilizes host lipids, such as cholesterol, for replication in infected cells. It has been suggested that cholesterol-rich membrane and lipid droplets (LD) play important roles in DENV entry and in the localization of viral capsid (C) proteins. The addition of PF-429242 showed suppression of viral propagation in all DENV serotypes^34,36^. Another study showed that in vitro selected PF-429242-DENV cultures have not replicated after five times sequential treatment with PF-429242. The intracellular levels of lipids reduced with the treatment, and viral propagation were not recovered by addition of exogenous lipids^34^. These observations potentially provide a possible approach for the effective treatment of YFV infection using PF-429242. In our assay the PF-429242 did not present activity against DENV2 (data not shown), and this divergence in activity levels is likely due to the assay specific methodology: PF-429242 was shown to impair dengue virion assembly and release, but did not interfere with viral replication and protein production; in the assay reported here, dengue viral levels are indirectly monitored through immunofluorescence of the E protein, and thus interference with protein levels is not expected with PF-429242.

Phospholipase C catalyse the hydrolysis of phosphatidylinositol (PI) into two intracellular messengers, diacylglycerol and inositol triphosphate (IP3), which mediate the activation of protein kinase C (PKC) and release of intracellular Ca^2+^, respectively. The compound U-73122 was originally described as a potent inhibitor of agonist-induced platelet aggregation by inhibiting PLC^56,57^. The analogue U-73343 differs from U-73122 by the substitution of a maleimide group for the less electrophilic succinide group, which resulted in loss of its PLC inhibitory ability, and therefore its analogue U-73343 is usually deployed as a negative control^58^. Chami et al. 2006^59^ reported calcium-signalling alterations related to the intracellular presence of complete virions or viral proteins. They showed that elevations of cytosolic calcium levels allowed an increased viral protein expression for HIV-1, viral replication for HBx, enterovirus 2B, HTLV-1 p12I, and EBV, viral maturation for rotavirus, viral release for enterovirus 2B and cell immortalization for EBV^39,59^. Another study demonstrated that the DENV2 infection causes an increased expression of the PLC, which cleaves the viral non-structural protein 1 (NS1) on the surfaces of cells infected with DENV. They suggested that the upregulation of PLC expression could induce an increase in the levels NS1 released from host cells and thereby contribute to the haemorrhagic symptoms characteristic of severe dengue^60^. Thus, while it is likely that U-73343 inhibits YFV and DENV infection through a similar or the same mechanism, and that it might involve PLC, it is not clear yet how it happens in molecular terms, and it would be interesting to determine U-73343 mechanism of action as it offers potential as a broad-spectrum antiflaviviral.

Flaviviruses exploit host cell pathways and machinery during their replicative life cycle, and the inhibition of these cellular processes might have an inhibitory effect on virus replication. Boldescu et al. 2017^32^ summarized the findings of numerous compounds with broad-spectrum antiviral activity that have already been identified by target-specific or phenotypic assays. Most broad-spectrum antiflaviviral compounds target viral protease or polymerase, host targets that are exploited during entry and replication, and proteins involved in nucleoside biosynthesis. Some anti-YFV compounds herein described were previously reported as having activity against several viruses and can serve as a potential starting point for the development of antiviral chemotherapy for yellow fever and other viruses.

## Acknowledgments

We would like to thank Prof. Dr. Paolo Marinho de Andrade Zanotto for kindly providing technical support. DRBP wishes to thank CNPq – Conselho Nacional para o Desenvolvimento Científico e Tecnológico for the PhD scholarship number 141603/2017-8.

